# Stress relaxation rates of myocardium from failing and non-failing hearts

**DOI:** 10.1101/2024.07.24.604845

**Authors:** Marissa Gionet-Gonzales, Gianna Gathman, Jonah Rosas, Kyle Y. Kunisaki, Dominique Gabriele P. Inocencio, Niki Hakami, Gregory N. Milburn, Angela A. Pitenis, Kenneth S. Campbell, Beth L. Pruitt, Ryan S. Stowers

## Abstract

The heart is a dynamic pump whose function is influenced by its mechanical properties. The viscoelastic properties of the heart, i.e. its ability to exhibit both elastic and viscous characteristics upon deformation, influence cardiac function. Viscoelastic properties change during heart failure (HF), but, direct measurements of failing and non-failing myocardial tissue stress relaxation under constant displacement are lacking. Further, how consequences of tissue remodeling, such as fibrosis and fat accumulation, alter the stress relaxation remains unknown. To address this gap, we conducted stress relaxation tests on porcine myocardial tissue to establish baseline properties of cardiac tissue. We found porcine myocardial tissue to be fast relaxing, characterized by stress relaxation tests on both a rheometer and microindenter. We then measured human left ventricle (LV) epicardium and endocardium human tissue from non-failing, ischemic HF, and non-ischemic HF patients by microindentation. We found that the ischemic HF had slower stress relaxation than non-failing endocardium; and that slower stress relaxing tissues were correlated with increased collagen deposition and increased α-smooth muscle actin (α-SMA) stress fibers, a marker of fibrosis and cardiac fibroblast activation, respectively. In the epicardium, we found that ischemic HF had faster stress relaxation than non-ischemic HF and non-failing; and that faster stress relaxation correlated with Oil Red O staining, a marker for adipose tissue. These data show that changes in stress relaxation vary across the different layers of the heart during ischemic vs. non-ischemic HF. These findings reveal how the viscoelasticity of the heart changes, which will lead to better modeling of cardiac mechanics for in vitro and in silico HF models.

## 1. Introduction

Viscoelasticity allows stress within the heart to dissipate over time. Previous studies based on in vivo imaging suggest that the viscoelasticity of the heart changes during HF^1, 2^. However, direct ex vivo viscoelastic measurements of human HF and non-failing tissue are lacking. Further, how tissue remodeling due to HF influences cardiac viscoelasticity remains unknown.

The heart’s functional performance is directly linked to its viscoelasticity. Viscoelasticity regulates cardiac wall compliance which in turn controls the amount of blood that can fill the chamber.^3^ The LV filling capacity is increased in faster relaxing cardiac tissue and decreased in slower relaxing cardiac tissue.^4^ If the heart does not relax fast enough for adequate filling, the subsequent decrease in LV output will cause an increase in end diastolic pressure. Increased diastolic pressure results in increased stress on the heart walls, leading to a variety of complications. In the short term, this increased pressure leads to circulatory congestion, causing edema;^5^ long term, high blood pressure can damage cells and lead to diastolic HF.^6^

#### Box: Viscoelasticity defined

##### Viscoelasticity

Viscoelastic materials exhibit both a viscous and elastic response when compressed by a force (**A**). The resulting material deformation can be defined as strain (**1**), and the force can be converted to stress (**2**), where L_0_ is the initial sample length, L_1_ is the sample length during indentation, and Δ is the difference between the two.

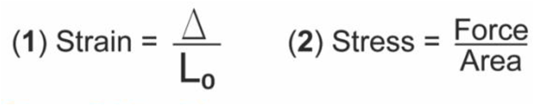

##### Stress Relaxation

In a stress relaxation test, a constant strain (**B**) is exerted on the material and the subsequent stress decay (**C**) is measured. We characterize the material relaxation by calculating the τ_1/2_. The τ_1/2_ indicates the time at which the stress reaches half its initial maximum value.

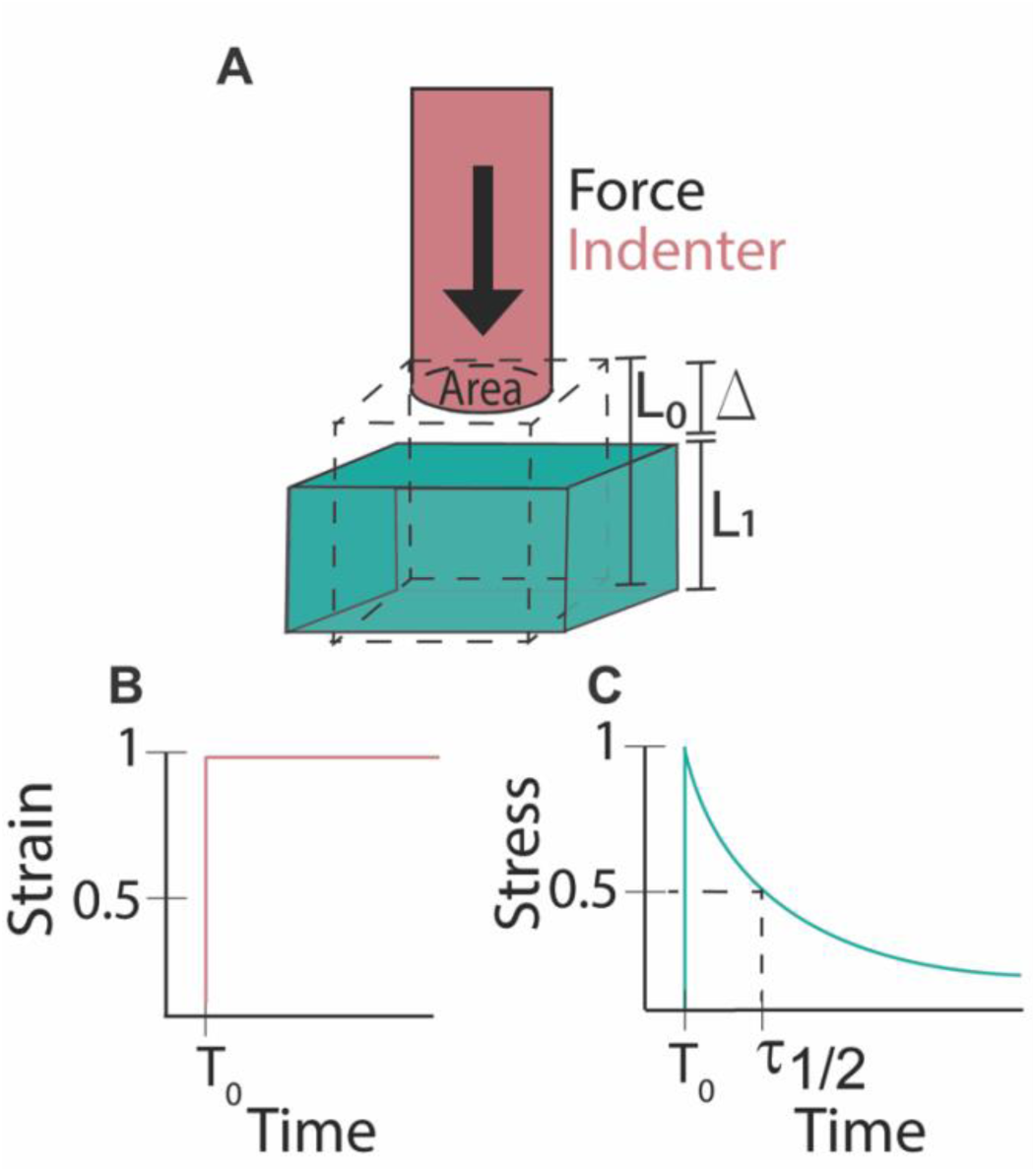

At the cellular level, viscoelasticity regulates cell proliferation^7^, spreading,^8^ differentiation^9^ and survival^10^ in a number of cell types, including fibroblasts,^11^ stem cells,^12, 13^ and cancer cells.^14^ To date, there has been limited investigation into how substrates with different relaxation rates influence cardiomyocytes (CMs). CMs are mechanosensitive, in vitro experiments show that they can modulate their morphology and function in response to stiffness^15–18^ and stretch.^19, 20^ Therefore, CMs are likely influenced by the viscoelastic cues of their substrate.

Viscoelastic substrates, including decellularized extracellular matrix (ECM),^21, 22^ and viscoelastic hydrogels, such as Matrigel, collagen and fibrin^23–25^, have been used for CM culture. However, these studies did not measure the viscoelasticity of their materials or investigate the effect of viscoelasticity independently of other parameters. Alternatively, biomaterials can be engineered to allow for tunability of viscoelasticity. Notably, one study found that CMs action potential duration increased after being crosslinked in a viscoelastic hydrogel compared to the cells before crosslinking.^26^ Another study showed increases in CM spreading, YAP localization, and sarcomere organization with increasing storage modulus, however they also saw an increase in their loss modulus and loss factor (tan δ), in their dynamic hydrogels.^17^ Since there were changes in both elastic and viscoelastic behavior within their hydrogels, the influence of viscoelasticity alone is hard to determine. Several other studies use viscoelastic engineered biomaterials for CMs, but are unable to verify that the viscoelasticity of their material is physiologically accurate due to the lack of reliable tissue measurements.^17, 27, 28^ Cell behavior can change substantially depending on the degree of substrate viscoelasticity^12^, motivating the need for cardiac viscoelastic measurements to aid in the development of cardiac biomaterials with physiologically relevant stress relaxation.

Computational models of the mechanical behavior of the heart would also benefit from experimental measurements of viscoelastic tissue mechanics.^29^ Several computational studies do not incorporate the effects of viscoelasticity.^30–32^ Studies that do investigate viscoelasticity, through incorporating experimental stress relaxation data, fail to agree on the role and importance of viscoelasticity in computational models. One reason for this disagreement is because it is unclear if the time-scale and magnitude of cardiac tissue viscoelasticity is relevant at a heartbeat scale; our work will help to answer this question.^33, 34^ One computational study concluded that viscoelasticity plays an important role in the cardiac relaxation phase, with increasing importance during disease conditions. ^35^ Through Finite Element Analysis, the authors found that their viscoelastic model had reduced stroke volume compared to an elastic model. Unfortunately, their model did not investigate how changing viscoelastic parameters, to mimic disease progression, influenced cardiac function. Our results address this gap in knowledge, by providing viscoelastic measurements from failing and non-failing human hearts.

To accomplish this, we conducted stress relaxation tests on porcine cardiac tissue and human non-failing cardiac tissues, as well as ischemic HF, and non-ischemic HF. We characterized fibrosis and adipose content within the endocardium (inner layer) and epicardium (outer layer), of the heart and correlated these with the relaxation rate. Our viscoelastic measurements will aid in the development of improved engineered biomaterials and computational modeling of the heart.

## 2. Materials and Methods

### 2.1 Porcine tissue acquisition and storage

We acquired porcine hearts to establish a baseline measurement of the relaxation of cardiac tissue. Porcine hearts are a useful approximation for human hearts, due to their similarity in structure, size and function.^36^ Frozen, formaldehyde-free porcine hearts were acquired from VWR (cat# 470330-138), or local markets (Santa Barbara Meat Co). Hearts were stored at −20°C, and thawed at 4°C for 12 h before mechanical testing was performed at room temperature. Once thawed, the LV was isolated using a scalpel. The LV wall was then cut with an 8 mm biopsy punch to produce a cylindrical core of tissue. The tissue core was then trisected into equal sections for mechanical testing. These sections were labeled as the inner layer (endocardium), middle layer (mid-myocardium), and outer layer (epicardium). Measuring the different layers independently is important since significant differences in the collagen content between the different layers can be indicative of heart disease.^37^

For mechanical measurements on decellularized tissue, we cut thawed porcine tissue into ∼15 × 5 mm sections of 2-3 mm thickness from the LV of a porcine heart. The sections were placed in 1% sodium dodecyl sulfate (SDS) on a plate shaker at 60 rpm at 4°C for 4 days. After this treatment the tissues were washed in 1X DPBS for 30 min to remove the remaining SDS. These decellularized tissues were then immediately subjected to mechanical testing.

### 2.2 Human tissue acquisition and storage

Human myocardium was procured from patients receiving cardiac transplants or from organ donors at the University of Kentucky. Patients, or their legally authorized representatives, provided informed consent and all procedures were approved by the University of Kentucky Institutional Review Board (protocol 46103). All the patients who received heart transplants had advanced HF. Hearts from organ donors were procured for research when the organ could not be used for transplant. Details of the procurement have been published.^38^ Samples of approximately 1 g were flash frozen in liquid nitrogen and shipped to UCSB on dry ice. The endocardium and epicardium were provided for each patient. Once received at UCSB, samples were stored at −80°C until day of testing. On the day of testing, samples were submerged in PBS to thaw for 5 min. Once thawed, a small flat section was cut off for testing, approximately 3-5 mm thick, with a surface of at least 3 × 3 mm.

### 2.3 Mechanical Characterization

Porcine cardiac tissue was measured using an ARES G2 rheometer (TA Instruments) with 8 mm parallel plates. First, an oscillatory shear strain test was conducted at 1% strain and an angular frequency of 6.28 rad/s to measure the storage and loss moduli of the tissue. Then, a stress relaxation test was performed, where an instantaneous 15% shear strain was applied to the sample and the resulting relaxation modulus was measured.

The viscoelasticity of human and porcine myocardium was measured using a custom-built microindenter.^39–43^ A 1.5 mm diameter quartz flat punch probe of 8 mm height was used for indentation. This allowed us to analyze smaller samples than would not have been possible using the rheometer. The lateral fixture had a spring constant of *K*_n_ = 437.8 μN/um, and the normal fixture had a spring constant of *K*_n_ = 184.6 μN/nm. The tissue was indented rapidly (2000 μm/s) to 10% of the sample thickness, and the indentation was held for at least 30 s, or until τ_1/2_, or half the initial maximum stress value, was reached. To confirm the relevance of our values for the heart, the percent of relaxation of the heart after one second, or the duration of an average heartbeat, was determined.

### 2.4 Histological Analysis

After mechanical testing, tissue samples were immediately embedded in optimal cutting temperature (OCT) compound, frozen on dry ice and sliced on a cryostat into 5 µm - 10 µm thick sections. For immunohistochemistry (IHC), sections were fixed in 4% paraformaldehyde (PFA) for 30 min. Samples were then permeabilized for 1 h in 0.1% Triton X-100 in phosphate-buffered saline (PBS). The samples were blocked for 3-4 h in a solution of 0.3% Tween20 and 2% bovine serum albumin (BSA) in PBS. Primary antibodies were diluted in blocking solution and incubated for 1 h at room temperature, or overnight at 4°C. Slides were washed several times with 0.1% Tween20 and 1% BSA in PBS, before incubation with the secondary antibodies diluted in blocking buffer for 1hr at room temperature. Slides were imaged using a Leica SP8 laser scanning confocal microscope with a 10x dry, 25x water-immersion objective and 63x oil-immersion objective for α-smooth muscle actin (α-SMA) fiber analysis.

For picrosirius red (PSR) staining, samples were fixed in 4% PFA for 30 min and stained in 0.1% PSR solution for 1 h at room temperature. Slides were then placed in a 0.01 N HCl solution for 2 min to wash away residual PSR. Slides were dehydrated in washes of increasing ethanol percentage and mounted with Permount mounting medium.

Oil Red O samples were fixed in 10% formalin for 30 min. The slides were then washed twice in running tap water, and dehydrated in 60% isopropanol for 2 min. They were then incubated with a 0.3% solution of Oil Red O for 2 min. Samples were then rinsed again with 60% isopropanol for 2 min, counterstained with hematoxylin and mounted with a glycerin solution.

### 2.5 Quantification of Histology

Large scan images were quantified for area of α-SMA and α-actinin. A threshold was applied to all channels, and particle analysis was performed using ImageJ. The area of all particles found on a large scan image was summed to acquire total area for a given channel. The area of α-SMA and α-actinin were normalized to DAPI area. For α-SMA stress fiber analysis, four representative 63x images were analyzed per slide via CT-FIRE^44^ with the threshold set to 1250. CT-FIRE gave us measurements for fiber width, length, and straightness.

Polarized light picrosirius red images were analyzed via CT-FIRE as well to characterize collagen fiber number, size and alignment. Brightness was adjusted in ImageJ on some images to enhance the number of fibers visible for quantification. Oil Red O images were quantified using a custom Python script (**Supplemental File 1**). The code used a saturation-based threshold to separate the sample from the background. The area occupied by the Oil Red O stain was then identified by pixels that fall under the HSV (hue, saturation, value) values for the colors red and pink. The rest of the sample (stained with hematoxylin) was identified by pixels with HSV corresponding to blues and purples. Using this script, the fat was quantified as the percentage of Oil Red O stained area over the total hematoxylin tissue area.

### 2.6 Collagen Quantification

Collagen content was determined using a hydroxyproline assay. Briefly, tissue samples were frozen, lyophilized, weighed, and then crushed using a pestle. The tissue was washed in PBS for 30 min, centrifuged, then dissolved in a 0.5 M acetic acid and 1 mg/mL pepsin solution, and stirred overnight at 4°C. Resultant tissue was centrifuged at 21,000 g for 30 min, and separated into pellet and supernatant. The supernatant was eluted to a new tube and placed on a hot plate at 105°C until only solid remained. Both tubes were mixed with 6 M HCl and heated to 105°C overnight. Sample solution was isolated, mixed with isopropanol, and incubated in 1:4 chloramine T and acetate citrate buffer for 10 min. Samples were then mixed with Ehrlich’s reagent and isopropanol (3:13) and incubated at 58°C for 30 min. The reaction was halted with ice and absorbance was read at 558 nm on Synergy H1 Multi-Mode Microplate Reader.

### 2.7 Statistical analysis

Normality of distribution was determined visually through Q-Q plots and histograms of the dataset. Human cardiac tissue τ_1/2_ data was non-normally distributed, as opposed to the porcine tissue which appeared normally distributed. Statistical analysis of human tissue was conducted via Kruskal-Wallis test for analysis between three or more groups, or Mann-Whitney test between two groups. All other data were statistically analyzed using one-way ANOVA or Student’s t-test.

## 3. Results

### 3.1.1 Porcine tissue was fast relaxing throughout all myocardial layers

To test our hypothesis that cardiac tissue is fast relaxing, the porcine left ventricle was isolated and characterized histologically and mechanically (**Figure 1**). We found aligned α-actinin positive CMs (**Figure 1B**), as well as a consistent concentration of approximately 3 μg collagen/mg tissue within each cardiac layer (**Figure 1C-D**). The storage modulus for all tissue layers was also very similar, around 2.5 kPa, with a slight increase in the average storage modulus for the epicardium at 2.9 kPa (**Figure 1E**). There was no significant difference in the loss modulus between layers, which were all around 0.6 kPa. Taken together, our measurements of the storage and loss moduli indicate limited differences in viscoelasticity between tissue layers, with only a slight difference between the endocardium and epicardium (**Figure 1E-F**). During stress relaxation tests, all layers of the tissue relaxed very rapidly, reaching τ_1/2_ in less than 6 s; corresponding well with our original hypothesis that cardiac tissue is fast relaxing. To determine the speed of relaxation, we quantified the τ_1/2_, or the time it takes for the stress to reach half of its maximum value. Interestingly, epicardium tissue relaxed roughly a full second slower than the other layers, at 5.9 s (**Figure 1G-H**).

**Figure 1.**
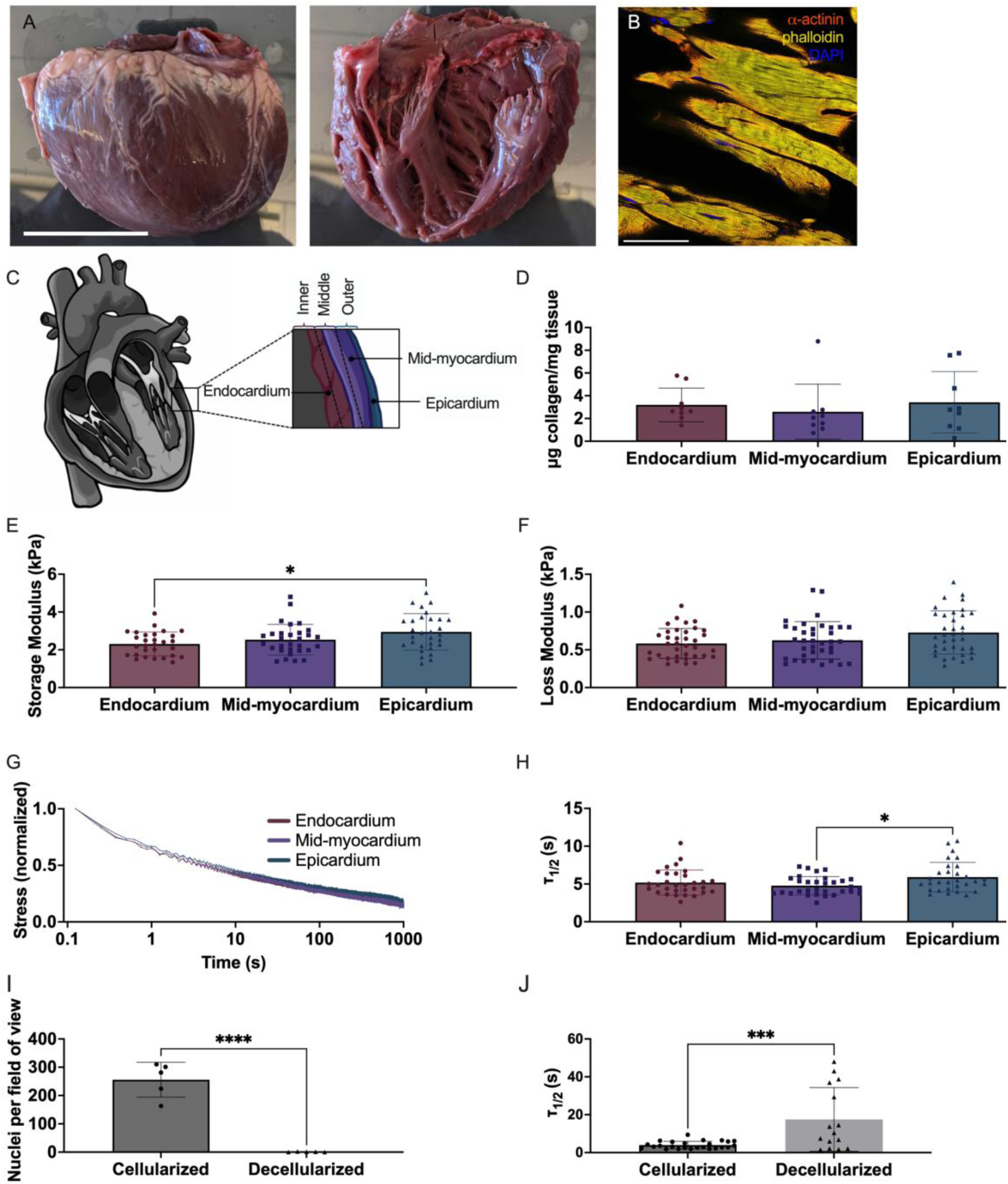
Porcine LV exhibits fast stress relaxation throughout all heart layers. **(A)** Gross image of excised porcine LV. **(B)** Immunohistochemistry image of porcine tissue stained for α-actinin (red). **(C)** Schematic of different layers of the heart. **(D)** Collagen content for each layer of the porcine LV was quantified via hydroxyproline assay. **(E)** Storage Moduli, **(F)** Loss Moduli, **(G)** Stress relaxation curves and **(H)** and τ_1/2_ were measured on the rheometer. **(I)** Porcine tissue was decellularized, verified by a depletion of DAPI. **(J)** Viscoelasticity of cellularized and decellularized tissue was measured on the microindenter. Only statistics with p value less than or equal to 0.05 are shown. *p ≤ 0.05, ***p ≤ 0.001, ****p ≤ 0.0001.

### 3.1.2 Decellularized porcine myocardium relaxed slower than cellularized porcine myocardium

We successfully decellularized the tissue, finding a significant depletion of nuclei in the decellularized tissue (**Error! Reference source not found.A-B**, **Figure 1I**). We measured the τ_1/2_ values for the cellularized tissue and decellularized tissue on the microindenter. We found that cellularized tissue relaxed significantly faster than decellularized, at an average τ_1/2_ of 3.9 s, while decellularized tissue had an average τ_1/2_ of 17.5 s (**Figure 1J**).

### 3.1.3 Stress relaxation tests performed via rheometry or microindentation yield similar trends in τ_1/2_ values

We compared τ_1/2_ values from the microindenter and rheometer, both in compression mode. There was a significant difference in the values, with the microindenter recording significantly faster τ_1/2_ values at 2.7 s compared to the rheometer at 5.5 s (Error! Reference source not found.). Although this was a significant discrepancy, the difference between the average of the two measurements was less than 3 s. Previous studies have shown that τ_1/2_ values of tissue can have a differences as large as 100 – 1000 s depending on tissue type^12^ and pathology.^14, 45^

### 3.2.1 Ischemic heart failure tissue exhibited slower stress relaxation in the endocardium and faster stress relaxation in the epicardium

Information on patients within each group is summarized in **Table 1**. For each patient, both the endocardium and epicardium of the LV were tested. The average stress relaxation curves for the endocardium indicated that non-failing tissues relaxed faster than non-ischemic and ischemic HF (**Figure 2A**). We used this data to quantify the time at which the stress reached half its maximum values, or τ_1/2,_ and found that non-failing was significantly faster than ischemic HF (**Figure 2C**). However, epicardium showed the opposite trend, with the ischemic HF group exhibiting the fastest stress relaxation (**Figure 2B**), and the τ_1/2_ data was in agreement, indicating ischemic HF was significantly faster than non-ischemic HF and non-failing (**Figure 2D**). We also compared the τ_1/2_ values of endocardium and epicardium between each group and found a significant difference in the endocardium and epicardium in the ischemic groups (Error! Reference source not found.). To justify why stress relaxation is important to consider for cardiac function, we analyzed the amount of relaxation that occurred within one second, the approximate time frame it takes to complete one human cardiac cycle. We found substantial stress relaxation occurred in just one second, with 35-40% of the initial stress relaxing on average in all groups, and significant decreases in percent of stress relaxation between non-failing and non-ischemic HF in the endocardium (Error! Reference source not found.).

**Table 1.** Patient demographics of indented cardiac tissue.

| Group | Age (years) | Sex (M/F) | Body mass index | Endo Average $\tau_{1/2}$ (s) | Epi Average $\tau_{1/2}$ (s) |
| --- | --- | --- | --- | --- | --- |
| Non-Failing (n=5) | 50.1 ± 9.7 | 3/2 | 30.9 ± 6.4 | 12.1 ± 19.9 | 14.6 ± 23.0 |
| Non-Ischemic HF(n=5) | 48.3 ± 9.6 | 3/2 | 29.5 ± 7.5 | 15.6 ± 27.1 | 11.3 ± 16.3 |
| Ischemic HF(n=5) | 50.3 ± 8.1 | 3/2 | 26.2 ± 2.7 | 15.8 ± 21.1 | 4.3 ± 3.8 |

**Figure 2.**
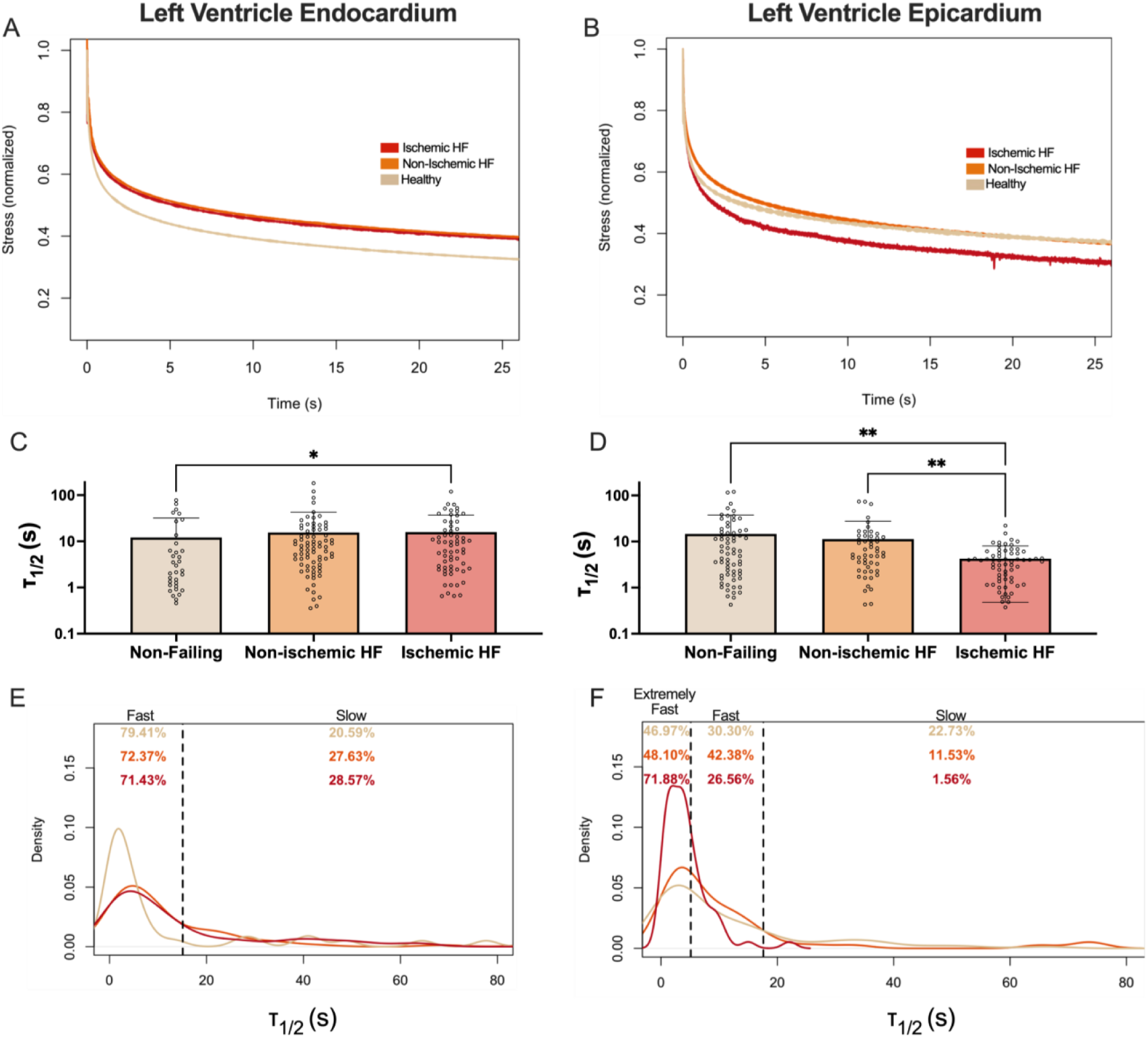
Stress relaxation changes during HF in the endocardium and epicardium. Average stress relaxation graphs for non-failing, ischemic and non-ischemic human tissue for **(A)** LV endocardium and **(B)** LV epicardium. Time at which stress reaches half the maximum value, or τ_1/2_, for **(C)** LV endocardium and **(D)** LV epicardium. Density graphs of τ_1/2_ values for **(E)** LV endocardium **(F)** LV epicardium. Dotted lines represent from left to right: reported τ_1/2_ values for fat (only in epicardium graph) and the average plus six standard deviations from the porcine tissue in figure 1. Percentages indicate percent of data located between dotted lines. Only statistics with p value less than or equal to 0.05 are shown. *p ≤ 0.05, **p ≤ 0.01.

### 3.2.2 Endocardial and epicardial tissues were categorized based on relaxation time into either slow, fast or extremely fast categories

We plotted the τ_1/2_ values as a density plot to visualize the spread of the data, and categorized the samples based on their viscoelasticity into either fast or slow relaxing tissues (**Figure 2E**). Moving forward, this separation of data based on stress relaxation allowed us to determine which biological differences were correlated to changes in stress relaxation.

We derived cutoff points using the average τ_1/2_ of the porcine data plus 6 standard deviations to capture the heterogeneity in the human data. The cutoff values for fast-relaxing tissue were 15.14s for the endocardium and 17.65s for the epicardium; samples with τ_1/2_ values greater than those cutoffs were considered slow-relaxing. In the endocardium data, we found that there was a distinct peak in fast τ_1/2_ values for the non-failing data that fell within the fast-relaxing regime, encompassing 79.41% of the data, while the ischemic and non-ischemic data had broader peaks with lower percentages within the fast-relaxing range (72.37% and 71.43% respectively) (**Figure 2E**). We hypothesized that fat was responsible for the fast stress relaxation of the epicardium. In order to explore this further, we introduced another cutoff point based on the τ_1/2_ of adipose tissue reported in the literature at 5.12s.^46^ This separation allowed us to determine if fat deposition correlated with fast stress relaxation. We termed the values faster than 5.12s extremely fast. The density plot of the epicardium showed that 71.88% of values for the ischemic HF group fall in the extremely fast category, compared with only 48.1% of non-ischemic HF and 46.97% of non-failing tissue samples. Additionally, the ischemic HF group had almost no samples (1.56%) within the slow relaxing regime, while non-failing tissue had the highest amount of slow stress relaxation values at 22.73%, followed by non-ischemic with 11.53% (**Figure 2F**). Throughout the rest of the paper, we will refer to the samples using the following categories for endocardium: fast (τ_1/2_ < 15.14s), and slow relaxing (τ_1/2_ > 15.14s); and for epicardium: extremely fast (τ_1/2_ < 5.12s), fast (5.12s<τ_1/2_ < 17.65s) and slow relaxing (τ_1/2_ > 17.65s).

### 3.3 α-actinin and α-SMA are increased in fast relaxing tissues

To correlate stress relaxation rates with tissue disease state, we stained indented tissue samples for α-actinin, a cytoskeletal actin-binding protein that is present in CMs, and α-SMA, a protein present in activated myofibroblasts. When analyzing the data based on the clinical categories, we saw no significant differences in α-actinin or α-SMA area (Error! Reference source not found.).

Qualitatively, we saw stark differences in α-actinin area between fast and slow relaxing groups in both endocardium and epicardium. In the endocardium, the fast-relaxing tissue had the most α-actinin, indicating the highest presence of CMs, while the slow relaxing tissue had far less α-actinin staining, and the α-actinin positive CMs present appeared smaller in length and diameter. However, there did not seem to be perceptible differences in α-SMA staining. (**Figure 3A**). The epicardium showed similar staining of α-actinin and α-SMA between tissues that were fast and slow relaxing. In the extremely fast relaxing group, we found a lack of staining for both α-SMA and α-actinin, indicating a lack of musculature (**Figure 3B**).

**Figure 3.**
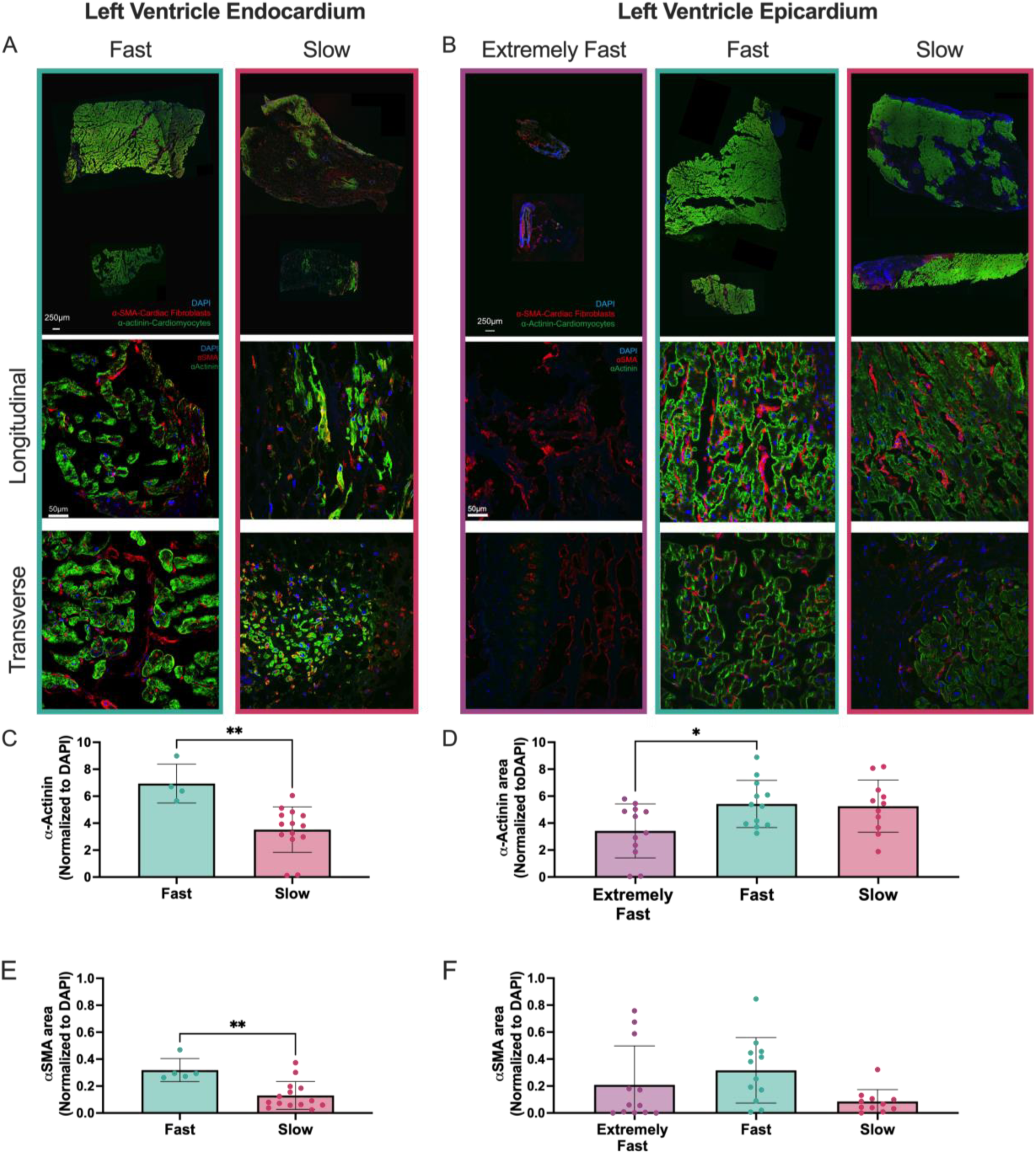
Fast relaxing tissue shows increased α-actinin and α-smooth muscle actin (α-SMA) in the endocardium and epicardium. **(A-B)** Representative large scans, and 25x images of longitudinal and transverse sections of endocardial and epicardial tissues within the extremely fast, fast, and slow relaxing categories. Images were stained with DAPI in blue, α-actinin in green, and α-SMA in red. Brightness and contrast on extremely fast epicardium large scan was increased more than others for better visualization. **(C-D)** Quantification of the area of α-actinin normalized to DAPI. (**E-F**) Quantification of the area of α-SMA normalized to DAPI. Only statistics with p value less than or equal to 0.05 are shown. *p ≤ 0.05, **p ≤ 0.01.

Upon quantifying the images, we found that α-actinin area was significantly greater in fast relaxing versus slow relaxing endocardium tissue (**Figure 3C**). The epicardium showed a significant increase in α-actinin stained area in fast relaxing tissue compared to extremely fast, and comparable amounts in slow relaxing (**Figure 3D**). We found significantly increased α-SMA staining in fast relaxing endocardial tissue compared to slow relaxing tissue (**Figure 3E**). In the epicardium, there was a trend of increasing α-SMA staining in fast relaxing tissue compared to slow, but it was non-significant (**Figure 3F**).

### 3.4 Slow relaxing endocardium contained larger α-SMA fibers

We were able to visualize α-SMA fibers in all stress relaxation groups in both the epicardium and endocardium (**Figure 4A-B**). α-SMA fibers are considered a more accurate representation of activated cardiac fibroblasts compared to α-SMA presence alone.^47^ Therefore, we quantified α-SMA fiber formation to determine if there was any difference in cardiac fibroblast activation between groups. We found a trend in the endocardium showing α-SMA stress fibers were longer (**Figure 4C**), significantly thicker (**Figure 4D**), significantly more aligned (**Figure 4G**) and on average present in greater numbers (**Figure 4H**) in slow relaxing tissues compared to fast relaxing tissue. In the epicardium we found no significant differences between the quantified α-SMA stress fibers with relationship to stress relaxation of the tissue (**Figure 4E-F, I-J**). However, we saw a trend of increased fiber number in the fast-relaxing group, indicating more activated fibroblasts compared to slow and extremely fast groups.

**Figure 4.**
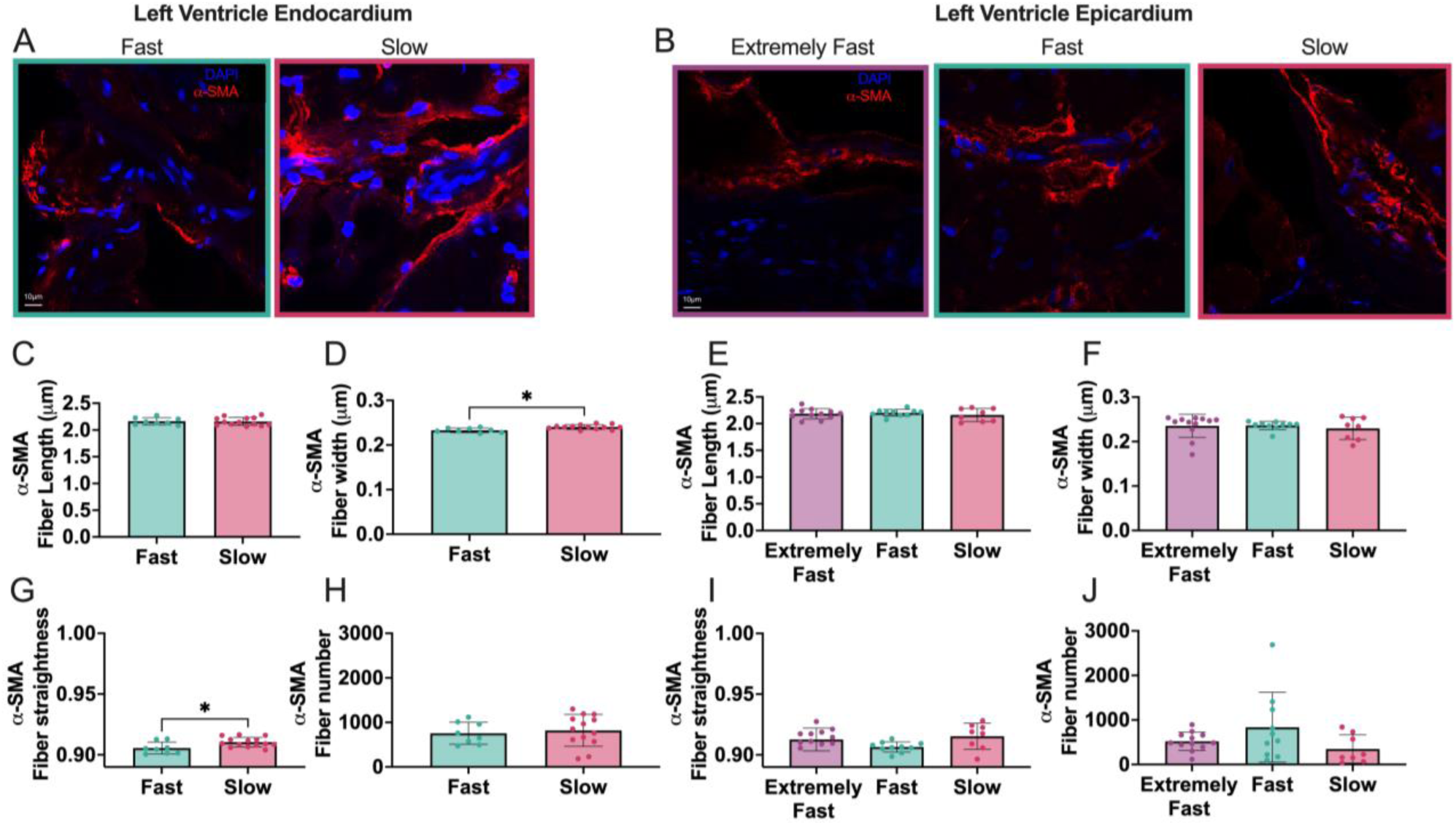
α-SMA stress fibers are thicker and straighter in slow relaxing endocardium tissue but show no significant trends in the epicardium. **(A-B)** 63x magnification images of α-SMA in extremely fast, fast and slow relaxing tissue of the endocardium and epicardium. Images were analyzed for α-SMA stress fiber **(C, E)** length, **(D, F)** width and **(G, H)** straightness. Graphs on the left represent endocardium, graphs on the right are epicardium. Only statistics with p value less than or equal to 0.05 are shown. *p ≤ 0.05.

### 3.5 Slow relaxing endocardium exhibited increased collagen staining and collagen fiber size

We stained the samples with picrosirius red and imaged them under polarized light to visualize the collagen fibers.^48^ Within the endocardium, we observed trends of increasing collagen amount in slower relaxing tissues. Qualitatively, the images in **Figure 5A** display more and brighter polarized collagen fibers in the slow relaxing group. In the epicardium, polarized light picrosirius red images revealed greater collagen abundance in the fast-relaxing group compared to extremely fast and slow relaxing tissues (**Figure 5B**). We found a significant increase in red pixels in these polarized images in slow relaxing endocardium tissue, denotating increased collagen amount. While in the epicardium the fast-relaxing tissues exhibited higher number of red pixels (**Figure 5C, E**). These results were in agreement with the hydroxyproline assay; which showed increasing collagen in slower relaxing endocardium and fast relaxing epicardium (**Figure 5D, F**).

**Figure 5.**
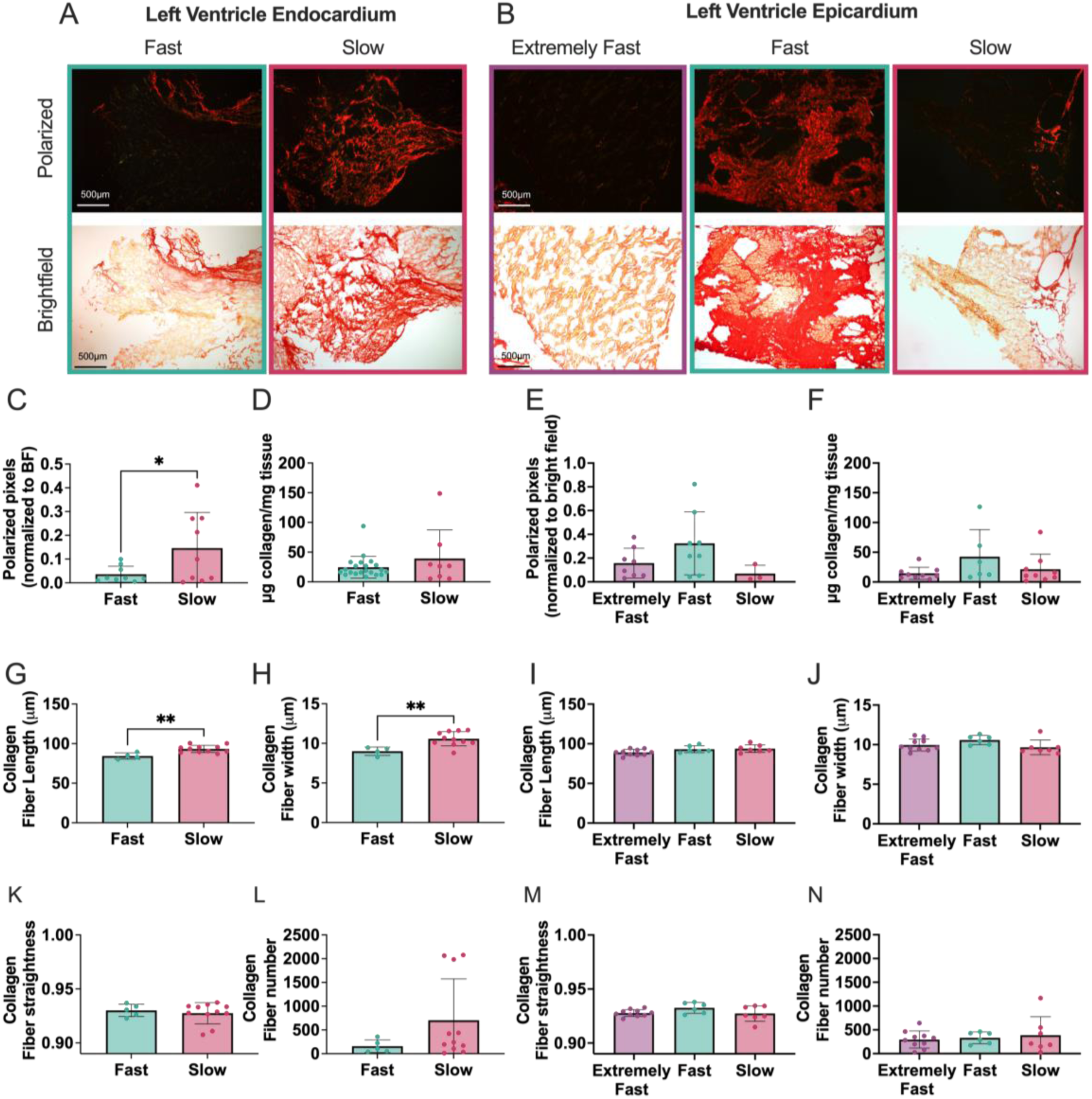
Collagen analysis shows increased fibrosis in slow relaxing endocardium but no significance in the epicardium. **(A-B)** Representative picrosirius red polarized and bright field images from endocardium and epicardium categorized as either extremely fast, fast or slow relaxing. Analysis of polarized pixels normalized to bright field area in **(C)** endocardium and **(E)** epicardium. **(D, F)** Hydroxyproline assay quantification of collagen concentration in tissue samples. Quantification of collagen fibers in picrosirius polarized light images, including **(G, I)** fiber length, **(H, J)** fiber width, **(K, M)** fiber straightness and **(LN)** fiber number. Only statistics with p value less than or equal to 0.05 are shown. *p ≤ 0.05, **p ≤ 0.01.

Through CT-FIRE analysis, we found that in the endocardium, collagen fibers were significantly longer and thicker in slower relaxing tissue than the faster relaxing tissues (**Figure 5G-H**). However, we found no significant differences in fiber straightness or fiber number between groups, although there was a distinct trend for increased fiber number in slow relaxing tissues (**Figure 5K-L**). In the epicardium, there was no significance in collagen fiber length, width, straightness or number (**Figure 5I-J, M-N**).

### 3.6 Epicardial tissue exhibited increased Oil Red O staining compared to the endocardium

We found that all epicardium tissues relaxed significantly faster than the endocardium; with epicardial tissue exhibiting an average τ_1/2_ of 10.1 s compared to an average τ_1/2_ of 15.0 s for endocardium (Error! Reference source not found.). We hypothesized that fat, which is known to be fast relaxing, was contributing to this difference. Although patient data was anonymized and de-identifiable, we knew which patients were diabetic and therefore likely had more epicardial fat.^49^ We analyzed our τ_1/2_ values based on patients with or without diabetes (Error! Reference source not found.**A**). Interestingly, in the endocardium we observed that patients with diabetes had significantly faster τ_1/2_, with a mean of 9.1 s, compared to those without diabetes, which exhibited a mean τ_1/2_ of 18.7 s. In the epicardium we observed similar trends, although they were not significant (Error! Reference source not found.**B**).

We then stained for lipids using Oil Red O to identify fat deposits in the tissue. Both fast and slow relaxing endocardium had limited Oil Red O staining (**Figure 6A**). However, the epicardium had substantially more in all groups, especially the extremely fast and fast relaxing tissues (**Figure 6B**). By quantifying the Oil Red O stain, we found that the aggregation of all epicardium groups had significantly higher amounts of fat than the aggregation of all epicardium groups (**Figure 6C**). When separating by stress relaxation times, we found the extremely fast, fast and slow epicardium had significantly higher Oil red O staining than the slow relaxing endocardium. There was also a trend of faster relaxing epicardium exhibiting higher average Oil Red O staining than slower relaxing epicardium, although this was not statistically significant (**Figure 6D**).

**Figure 6.**
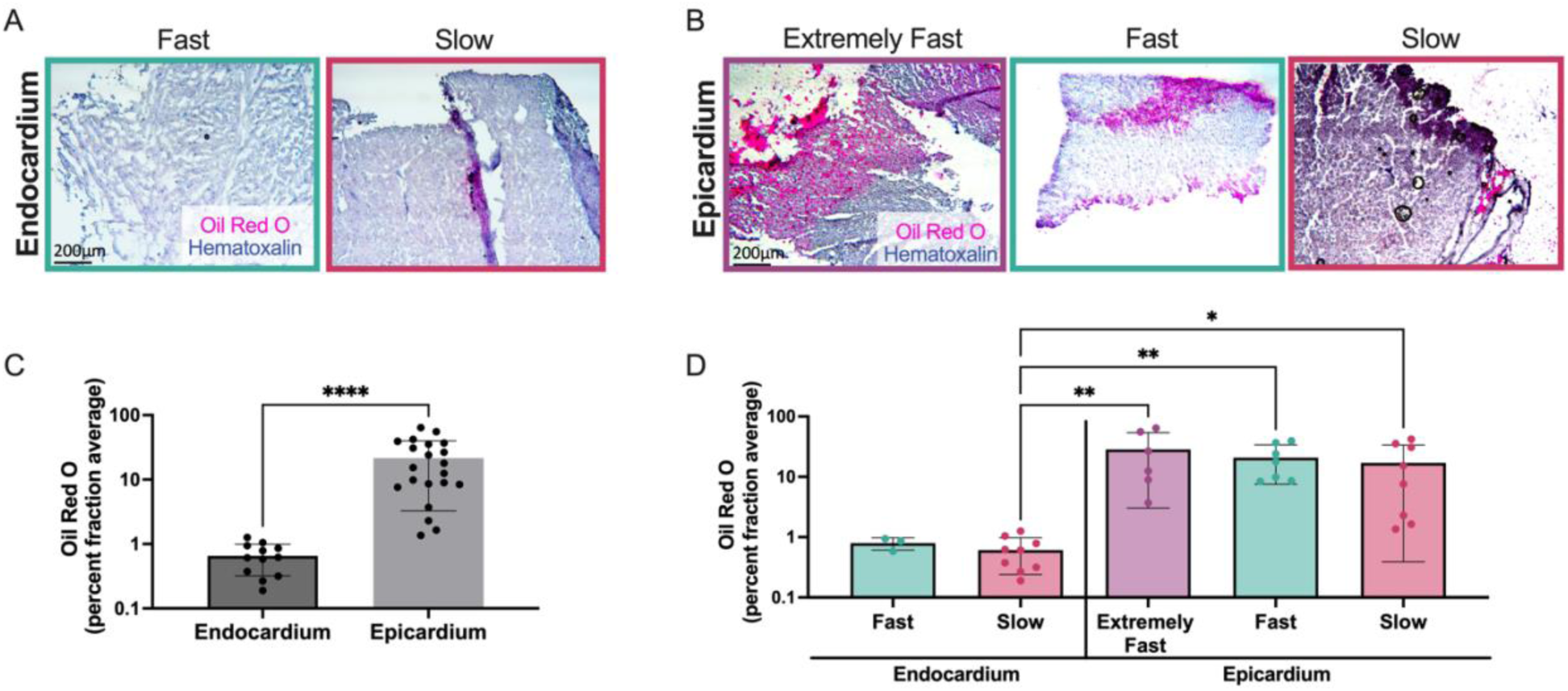
Oil Red O staining is upregulated in extremely fast relaxing epicardium. **(A-B)** Representative Oil red O stain from endocardium (left) and epicardium (right). **(C)** Quantification of Oil Red O stain for endocardium and epicardium. **(D)** Quantification of Oil red O separated based on stress relaxation speed for both endocardium and epicardium. Only statistics with p value less than or equal to 0.05 are shown. *p ≤ 0.05, **p ≤ 0.01, ****p ≤ 0.0001.

## 4. Discussion

We have shown, through mechanical tests that viscoelasticity changes with respect to disease state within the heart. We then used histological and biochemical analysis of collagen and fat to correlate local tissue composition and structure with stress relaxation. Although there have been notable viscoelastic measurements of cardiac tissue previously, both directly^34, 50–54^ and indirectly through imaging modalities^55, 56^, these measurements are difficult to translate to in vitro and in silico models. Several cardiac tissue measurements are conducted on custom-built instruments, making it difficult to reproduce viscoelastic measurements on other instruments. These tests also vary in the type of test performed, including stress relaxation, creep or oscillating stress, and if the force is applied in compression, tension, or shear. This concern is best demonstrated by a prior study comparing different mechanical measurement methods for the same cell source, resulting in material parameter estimations that were orders of magnitude apart across methods.^57^ Although we found it necessary to use a specialized instrument for our studies, due to small specimen size, we validated our measurements on a commercially available rheometer. We further conducted a simple compression stress relaxation test at 10% strain that can be replicated by several commercially available instruments including a rheometer, Instron or dynamic mechanical analyzer. Sample species, myopathy, storage and preparation can also lead to variation in reported results. Our HF and non-failing cardiac samples were acquired from a reputable human biobank, increasing the likelihood that our samples would produce relevant results.^38^

Porcine hearts were highly viscoelastic (∼5 s), relaxing much faster than other soft tissues (∼100 s – 1000 s).^12, 58, 59^ When decellularized, the tissue became slower relaxing (17.5 s), suggesting that both cells and ECM contribute to the viscoelastic response. Our data is in agreement with previous studies that have shown that microtubules and sarcomeres within CMs^29, 60–62^ as well as cardiac ECM^63^ are viscoelastic. This data counters studies that determined the characteristic relaxation time of the heart to be 600s, based on a fit to a Maxwell element model, which is far too long to be relevant at a heartbeat scale.^34^ Contrary to their findings, we found that even within 1s, the tissue relaxes around 40%, which likely influences cardiac function (Supplemental Figure 4).

We found faster relaxing tissue had more α-SMA, which was in agreement with in vitro studies showing fast relaxing hydrogels induce increased α-SMA expression in myofibroblasts.^64^ However, activated cardiac fibroblasts are known to not only express α-SMA, but form α-SMA stress fibers when fully differentiated.^47^ We found a trend of increasing α-SMA stress fiber number in slower relaxing tissue, indicating more activated fibroblasts. We also found significant increases in fiber width and alignment in slow relaxing tissues. Our reported differences in the stress fibers are a unique finding, since there is little quantitative data within the literature on stress fiber size and alignment in cardiac tissue. Previous in vitro studies have only qualitatively noted that fibroblasts stress fiber thickness changes due to substrate properties, with fibroblasts on silicone exhibiting thinner fibers than those on plastic.^65^

In the epicardium, fat serves a number of vital functions. However, excessive accumulation of epicardial fat can contribute to HF.^66 49^ We found that increases in fat and decreases in CMs led to faster tissue stress relaxation, likely contributing to further dysregulation of heart function. It is known that different cell types can exhibit distinct mechanical properties, therefore it is likely that this change in cell populations, regardless of matrix remodeling, has a direct impact on the stress relaxation rate.^67^

A limitation of this work is that we could only mechanically characterize the heart tissue in the passive state, since our tests were performed ex vivo, and thus our measurements are most relevant to diastole. During contraction, the activation of cross bridges within the CMs changes the tissues stiffness, and likely changes tissue viscoelasticity.^68^ Additionally, the heterogeneous nature of cardiac tissue was an expected challenge within this study. For example, scarring associated with HF is not usually present throughout all of the heart.^69^ To address tissue heterogeneity, we compared the biochemical and histological data from each tissue sample based on its τ_1/2_ value. This analysis yielded significant differences between fast and slow relaxing tissues, compared to classifying the data based on the clinical categorization which showed no significance (**Supplemental Figure 5**). Patient-to-patient variability was another challenge, as we only had access to 5 patients in each group. Even with our limited sample size, we showed that comorbidities can influence cardiac viscoelasticity. Our data indicated that diabetic patients, across HF and non-HF groups, had faster relaxing epicardium (**Error! Reference source not found.**). Investigation into what other patient characteristics effect viscoelasticity could lead to interesting insights into cardiac disease.

## 5. Conclusion

We have shown that both human and porcine heart is fast relaxing through direct stress relaxation tests on ex vivo tissue. Our studies further conclude that, across all groups, the endocardium becomes slower relaxing during HF due to fibrosis. The epicardium, however, shows a reverse trend; becoming faster relaxing during HF due to increased fat deposition. These results have important implications for computational and biomaterial cardiac disease modeling.

