## Supplemental Figures for "Stress relaxation rates of myocardium from failing and non-failing hearts"

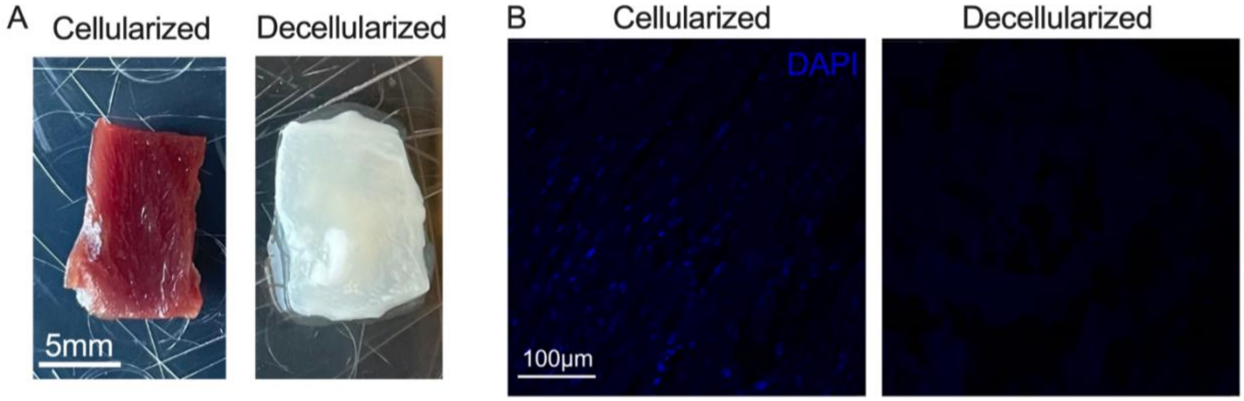


**Supplemental Figure 1. Decellularization was confirmed through a decrease in DAPI staining. (A)** Porcine tissue was decellularized, **(B-C)** verified by a depletion of DAPI staining. **(D)** $\tau_{1/2}$ values of cellularized and decellularized porcine tissue.

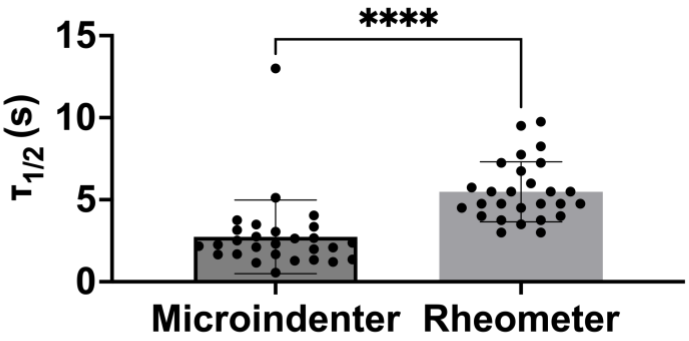


**Supplemental Figure 2. Porcine LV exhibited fast stress relaxation as measured on the rheometer and microindenter.** Measurement of $\tau_{1/2}$ of porcine LV myocardium from the microindenter and rheometer in compression mode. Only statistics with p value less than or equal to 0.05 are shown. ****$p \leq 0.0001$.

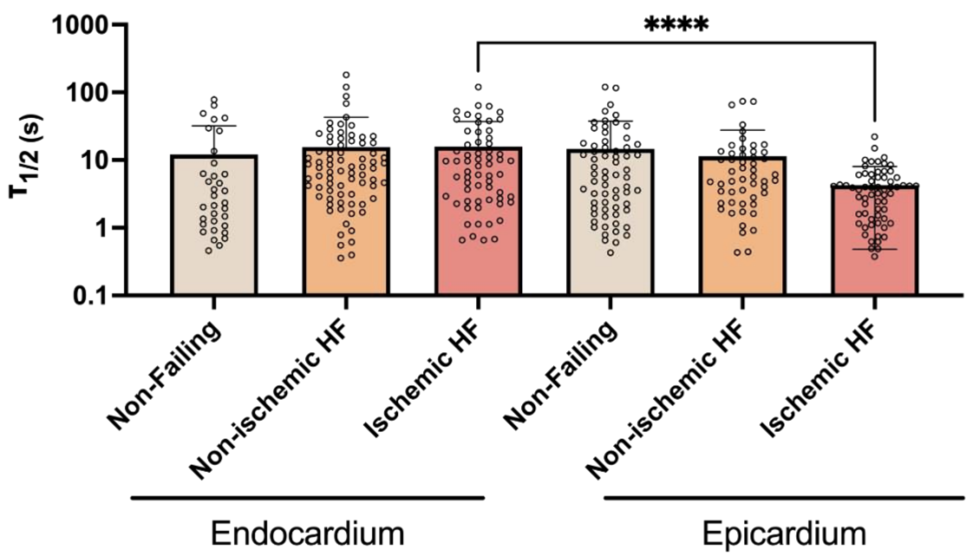


**Supplemental Figure 3. Ischemic HF group showed significant differences in $\tau_{1/2}$ between endocardium and epicardium.** Statistical comparison between endocardium and epicardium. Only statistics with p value less than or equal to 0.05 are shown. ****$p \leq 0.0001$.

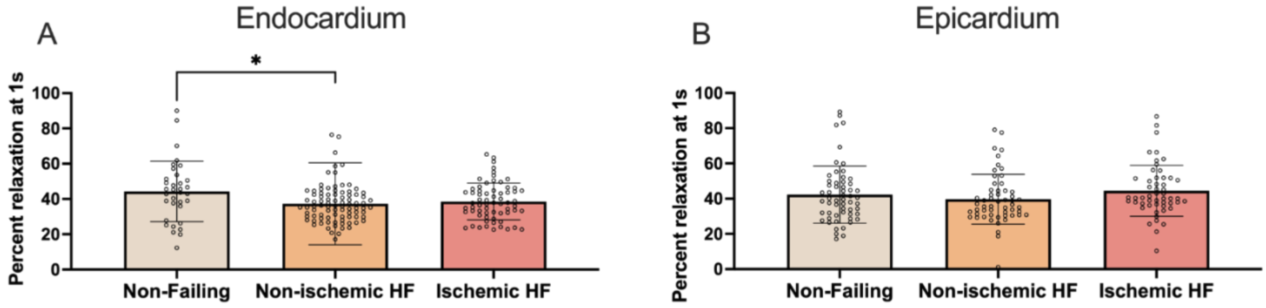


**Supplemental Figure 4. After 1s significant percent stress relaxation occurs in the epicardium and endocardium.** Percent stress relaxation occurring after 1s in the **(A)** endocardium and **(B)** epicardium. Only statistics with p value less than or equal to 0.05 are shown. *$p \leq 0.05$.

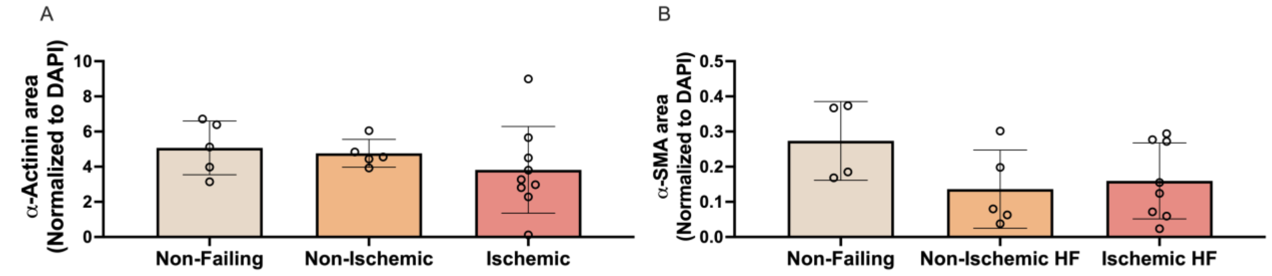


**Supplemental Figure 5. α-actinin and α-SMA staining quantification for the endocardium separated by clinical labels show no significance.** Quantification of the area of **(A)** α-actinin and **(B)** α-SMA normalized to DAPI for endocardium. Only statistics with p value less than or equal to 0.05 are shown.

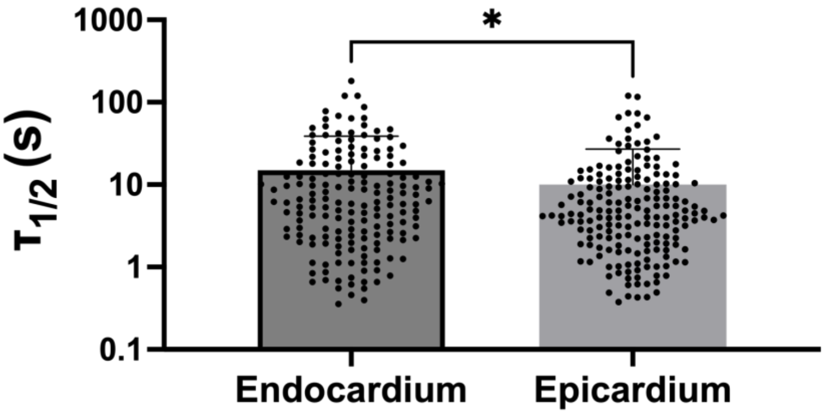


**Supplemental Figure 6. The endocardium shows significantly higher average $\tau_{1/2}$ values compared to the epicardium.** Measurement of $\tau_{1/2}$ between all human endocardium and epicardium samples. Only statistics with p value less than or equal to 0.05 are shown. $^*p \leq 0.05$.

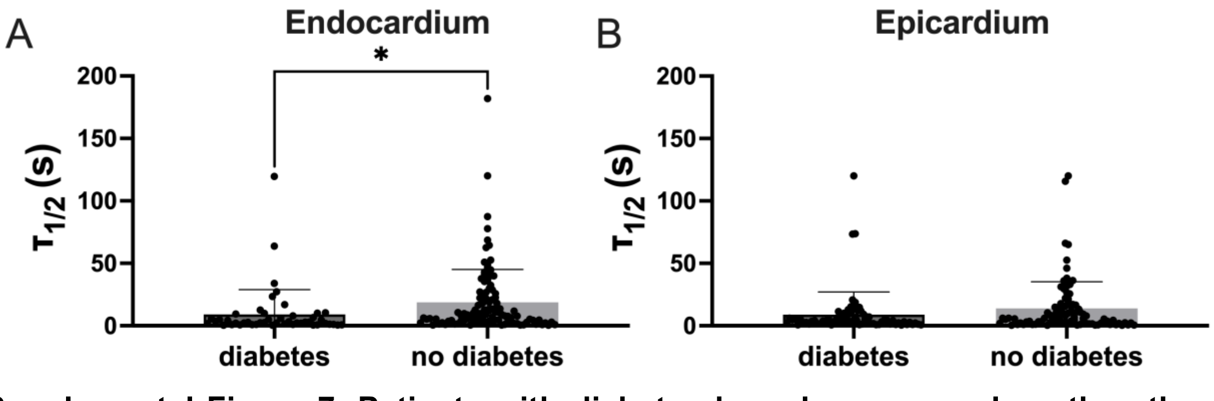


**Supplemental Figure 7. Patients with diabetes have lower $\tau_{1/2}$ values than those without diabetes.** $\tau_{1/2}$ values for **(A)** endocardium and **(B)** epicardium separated by patients who have or do not have diabetes. Only statistics with p value less than or equal to 0.05 are shown. $^*p \leq 0.05$.
