## Supplemental File.1 for "Stress relaxation rates of myocardium from failing and non-failing hearts"

ored\_o\_analysis


In [1]:

```
import numpy as np
import cv2 as cv
import pandas as pd
import sys, glob, os, fnmatch
from matplotlib import pyplot as plt
```

In [2]:

```
def load_images(folder):
    images = []
    for name in os.listdir(folder):
        img = cv.imread(os.path.join(folder, name))
        if img is not None:
            images.append(img)
    return images
    
def resizing(img, scale):
    # scale_percent = scale
    width = int(img.shape[1]*scale/100)
    height = int(img.shape[0]*scale/100)
    dim = (width, height)
    resized = cv.resize(img, dim, interpolation = cv.INTER_AREA)
    return resized

#Determining tissue area
def color_detect(img):
    sat = cv.cvtColor(img, cv.COLOR_BGR2HSV)[...,1]
    value = cv.cvtColor(img, cv.COLOR_BGR2HSV)[...,2]
    _, mask = cv.threshold(sat,100,255,cv.THRESH_BINARY)
    _, mask2 = cv.threshold(value,200,255,cv.THRESH_BINARY)
    masks = mask + mask2
    color_out = cv.bitwise_and(img,img,mask=mask)
    background = np.where(color_out > 100, 255, 0).astype(np.uint8)
    # background = cv.cvtColor(background, cv.COLOR_GRAY2BGR)
    # background = cv.cvtColor(background, cv.COLOR_BGR2HSV)

    col0r_out = color_out + background
    tissue_pix = cv.countNonZero(mask)
        
    return tissue_pix, color_out

#Determining Oil Red O area
def oil_red(img):
    hsv = cv.cvtColor(img, cv.COLOR_BGR2HSV)

    red_low_1 = np.array([0,70,30])
    red_upp_1 = np.array([2,255,255])
    mask1 = cv.inRange(hsv, red_low_1, red_upp_1)

    red_low_2 = np.array([140,70,30])
    red_upp_2 = np.array([179,255,255])
    mask2 = cv.inRange(hsv, red_low_2, red_upp_2)

    redmask = mask1 + mask2

    red_out = cv.bitwise_and(img,img,mask=redmask)
    red_ratio = cv.countNonZero(redmask)

    return red_out, red_ratio

    #blue mask 

#Determining hematoxylin area
def blue_stain(img):
    hsv = cv.cvtColor(img, cv.COLOR_BGR2HSV)
    # blue mask
    lower = np.array([110,35,0], np.uint8)
    upper = np.array([140,255,255], np.uint8)
    # lower = np.array([60, 35, 140])
    # upper = np.array([180, 255, 255])
    blue_mask = cv.inRange(hsv, lower, upper)
    blue_out = cv.bitwise_and(img,img,mask=blue_mask)
    blue_ratio = cv.countNonZero(blue_mask)

    return blue_out, blue_ratio
```

In [3]:

```
samplename = 'S4'
images = load_images(samplename)

num_images = len(images)
```

In [4]:

```
tissue_pix = []
red_pix = []
blu_pix = []

for i in images:
    tissue_area, color_out = color_detect(i)
    red_out, red_count = oil_red(i)
    blu_out, blu_count = blue_stain(i)
    red_concat = cv.hconcat([i, red_out])
    blu_concat = cv.hconcat([i, blu_out])
    concat = cv.vconcat([red_concat, blu_concat])
    concat = resizing(concat, 50)
    tis_pix = red_count+blu_count
    tissue_pix.append(tis_pix)
    red_pix.append(red_count)
    blu_pix.append(blu_count)
    concat = cv.cvtColor(concat, cv.COLOR_BGR2RGB)
    plt.imshow(concat)
    plt.title('Oil Red O')
    plt.xticks([]), plt.yticks([]) 
    plt.show()

#Oil Red O stain / total tissue area
red_tissue_ratio = [x/y for x,y in zip(red_pix, tissue_pix)]
#Hematoxylin stain / total tissue area
blu_tissue_ratio = [x/y for x,y in zip(blu_pix, tissue_pix)]
#Oil Red O / hematoxylin ratio
red_blu_ratio = [x/y for x,y in zip(red_pix, blu_pix)]

#Creating dataframe 
dict = {'Sample Number': list(range(1, len(images)+1)),
        'Red Pixels': red_pix,
        'Blue Pixels': blu_pix,
        'Red/Blue Fraction': red_blu_ratio,
        'Red/Total Fraction': red_tissue_ratio
       }

df = pd.DataFrame(dict)
cv.destroyAllWindows()
```

In [6]:

```
df.head()
```

Out[6]:

|  | Sample Number | Red Pixels | Blue Pixels | Red/Blue Fraction | Red/Total Fraction |
| --- | --- | --- | --- | --- | --- |
| 0 | 1 | 366106 | 355390 | 1.030153 | 0.507426 |

In [22]:

```
#Saving data table to .csv
df.to_csv('Epi Data/Oil Red O/61BAC_S4.csv', index=False)
```
